# TriMem: A Parallelized Hybrid Monte Carlo Software for Efficient Simulations of Lipid Membranes

**DOI:** 10.1101/2022.05.25.493239

**Authors:** Marc Siggel, Sebastian Kehl, Klaus Reuter, Jürgen Köfinger, Gerhard Hummer

**Affiliations:** European Molecular Biology Laboratory Hamburg, Notkestraße 85, 22607 Hamburg, Germany; Centre for Structural Systems Biology (CSSB), Notkestraße 85, 22607 Hamburg, Germany; Department of Theoretical Biophysics, Max Planck Institute of Biophysics, Max-von-Laue-Straße 3, 60438 Frankfurt am Main, Germany; Max Planck Computing and Data Facility, Gießenbachstraße 2, 85748 Garching, Germany; Institute of Biophysics, Goethe University Frankfurt, 60438 Frankfurt am Main, Germany

## Abstract

Lipid membranes are integral building blocks of living cells and perform a multitude of biological functions. Currently, molecular simulations of cellular-scale membrane structures at atomic resolution are nearly impossible, due to their size, complexity, and the large times-scales required. Instead, elastic membrane models are used to simulate membrane topologies and transitions between them, and to infer their properties and functions. Unfortunately, efficiently parallelized open-source simulation code to do so has been lacking. Here, we present TriMem, a parallel hybrid Monte Carlo simulation engine for triangulated lipid membranes. The kernels are efficiently coded in C++ and wrapped with Python for ease-of-use. The parallel implementation of the energy and gradient calculations and of Monte Carlo flip moves of edges in the triangulated membrane enable us to simulate also large and highly curved sub-cellular structures. For validation, we reproduce phase diagrams of vesicles with varying surface-to-volume ratios and area difference. The software can tackle a range of membrane remodelling processes on sub-cellular and cellular scales. Additionally, extensive documentation make the software accessible to the broad biophysics and computational cell biology communities.

## I. INTRODUCTION

Living cells are bounded by lipid membranes, and the interior of eukaryotic cells is filled with membranous organelles. Cellular membrane structures are highly dynamic, strongly curved, and branched^1,2^. Membranes are flexible and behave like two-dimensional fluids. Super-resolution microscopy and cryo-electron tomography (cryo-ET) have given unprecedented insights into the spatial organization of cellular membranes and their associated protein structures^3^. However, a detailed and comprehensive biophysical understanding of the mechanisms underlying organellar membrane reshaping in the cellular context remains elusive.

Molecular dynamics simulations are increasingly becoming state of the art for understanding membrane biophysics in the cellular context^4–7^. Multi-scale approaches combining all-atom, coarse-grained and meso-scale methods are used to tackle various aspects of membrane remodeling^8,9^. All-atom models are used to gain detailed insight into protein-lipid interactions and protein-protein interactions inside membranes but are limited in size and time scale^9^. Coarse-grained particle-based simulations approaches make it possible to study large membrane-associated complexes and complex membrane shape changes^9–12^. However, simulations of large cell-scale membrane systems are currently impossible with molecular models at atomic or near-atomic resolution. Computational resources limit the time and length scales that can be studied with particle-based approaches. Therefore, sub-cellular and cell-scale remodelling processes of organelles, vesicles, and cells are commonly studied by using meso-scale models and continuum approaches, primarily membrane-elastic theory^13,14^.

Dynamic triangulated surfaces (DTSs) have emerged as a useful meso-scale model to solve the Helfrich Hamiltonian numerically and study large-scale membrane-shaping processes. Initially, DTSs were used to study shaping and properties of fluctuating giant unilamellar vesicle (GUV)^15–18^. The discretization of membranes makes it possible to sample non-axisymmetric shapes^17^ beyond symmetric arc length parameterizations^19,20^. This method has been applied to a number of biological processes with increasing complexity including nano-particle wrapping^21,22^, membrane tubulation^23^, formation of autophagic vesicles^24^, formation of Golgi stacks^25^, and protein-induced membrane budding^26,27^.

Here, we present TriMem, an open-source software package for efficient simulation and optimization of DTSs using a parallelized hybrid Monte Carlo (MC) approach. This approach overcomes limitations of existing serial implementations. In such non-parallel approaches, the number of vertices of the triangulated membrane representation is severely limited to keep computational times manageable. Yet, large numbers of vertices are needed to describe highly curved membranes, which are ubiquitous in cellular organelles such as mitochondria or the tubular ER. As a further challenge, the commonly used random single-particle MC moves are inefficient with respect to sampling. Importantly, only few code bases for these types of simulations have been made freely available to the broader community despite the large amount of scientific research in this area, and none of them are parallelized yet^28,29^.

This paper is structured as follows. First, we briefly recapitulate the discretized version of Helfrich theory and the Helfrich Hamiltonian used throughout this work (section II). We then introduce the TriMem software, including its implementation strategy and algorithms (section III). We present benchmark results on timings for varying mesh sizes and compare the performance to a serial single-core implementation (section III D). We validate the software by reproducing well established phase diagrams of vesicle shapes (section IV). Finally, we provide a comprehensive outlook on the expected impact of our software and on future developments (section VI)

## II. MEMBRANE MODEL

Membrane-elastic theory describes membranes as 2D surfaces with fluid properties embedded in 3D space. In its simplest form, the bending free energy of a membrane can be written in terms of the so-called Helfrich Hamiltonian^13,14^,

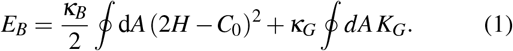

with *H* = 0.5(*c*_1_ + *c*_2_) the mean curvature and *K*_*G*_ = *c*_1_*c*_2_ the Gaussian curvature, where *c*_1_ and *c*_2_ are the principal curvatures. *C*_0_ signifies the intrinsic curvature of the membrane which is modulated by a variety of factors, including proteins, lipid composition and membrane asymmetry. *κ*_*B*_ and *κ*_*G*_ are the bending rigidity and Gaussian bending modulus, respectively, and describe the elastic properties of the studied membrane. The second term, the Gaussian curvature, can usually be neglected because it is constant in the absence of changes of the topology according to the Gauss-Bonnet theorem. Multiple extensions and variations of the Helfrich Hamiltonian have been introduced to include a broad spectrum of external factors such as area difference, protein inclusions, and osmotic pressure. Variational minimization of the Helfrich Hamiltonian with respect to the membrane shape leads to fourth-order differential equations, which have been solved analytically only for a limited number of cases of high symmetry^30,31^. However, numerical solutions are achievable.

To make simulations of membrane shapes computationally tractable and amenable to integration and MC sampling, the surface is discretized as a triangular mesh, which can in principle represent arbitrary surface shapes. The Hamiltonian of this discretized system (*H*_tot_) is more complex than the original Helfrich Hamiltonian. We decompose the energy as

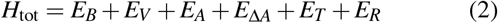

where *E*_*B*_ is the bending energy, *E*_*V*_ the volume energy to maintain the total internal volume, *E*_*A*_ the area energy to maintain the total surface area, *E*_Δ*A*_ the area-difference energy (ADE), *E*_*T*_ the tethering potential, and *E*_*R*_ the repulsive potential. In the following, we explain how we calculate these energies from triangulated surfaces.

To introduce the energy terms, we first have to define the DTS. A closed surface system consists of *N*_*T*_ = 2(*N*_*V*_ − 2) triangles, with *N*_*V*_ vertices connected by 3(*N*_*V*_ − 2) tethers. To this end we introduce a triangulation *𝒥*:= (**x**, *F*) as a tuple of vertex positions 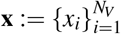 (with a single vertex *x*_*i*_ ∈ ℝ^3^) and triangles 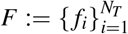. Thereby a single oriented triangle *f*_*l*_: = {(*i, j,k*)|*i, j,k* ∈ [1, *N*_*V*_], *i* ≠ *j* ≠ *k*} is given by an ordered 3-tuple indexing into the vertices **x**. For the convenience of notation, we define the vertex-vertex connectivity 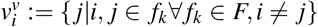, the vertex-face connectivity 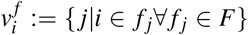, the set of edges *E* := {(*i, j*)|*i, j* ∈ *f*_*k*_∀ *f*_*k*_ ∈ *F, I* ≠ *j*} and the edge-face connectivity 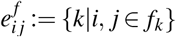

We discretize the calculation of the Helfrich Hamiltonian *H*_tot_ on the DTS using a vertex-averaged formulation as previously described extensively in ref. 17. The total bending energy *E*_*b*_ is given by the sum over all vertices of the bending energy per vertex

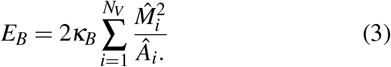

The area *Â*_*i*_ per vertex *i* is calculated as

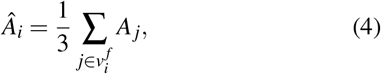

where *A*_*j*_ is the area of the single triangle *f* _*j*_. The average mean curvature 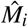 associated with vertex *i* can be calculated as sum over all adjacent edges

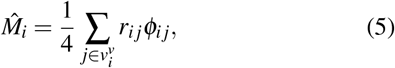

where *r*_*i j*_ = ‖*x*_*j*_ − *x*_*i*_‖ and *ϕ*_*ij*_ is the angle between the oriented normal vectors *n*_*a*_, *n*_*b*_ of the triangles in 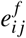, i.e., adjacent to the edge (*i, j*), calculated as cos(*ϕ*_*i j*_) = *n*_*a*_ · *n*_*b*_.

The total mean curvature of the system is then computed by

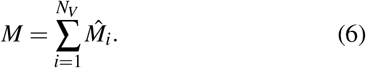

The energies for the volume and area constraint are given by *E*_*V*_ and *E*_*A*_, respectively, and can be written as simple harmonic potentials

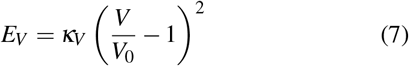

and

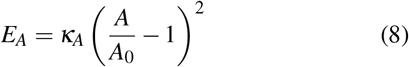

with *κ*_*V*_, *κ*_*A*_ being the coupling constants, respectively. The corresponding reference values are given by *V*_0_ and *A*_0_. *κ*_*V*_ and *κ*_*A*_ are usually chosen several orders of magnitude larger than *κ*_*B*_ to avoid non-physical area and volume fluctuations. The total area *A* is calculated as

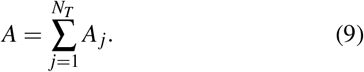

Also the volume *V* can be calculated as a discrete sum,

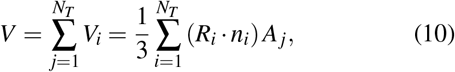

where *V*_*i*_ is the signed sub-volume of a single triangle, *R*_*i*_ the position vector, and *n*_*i*_ the unit normal vector.

The ADE, *E*_Δ*A*_, is given by

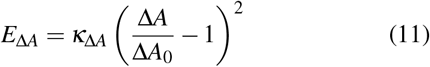

The area difference Δ*A* of a bilayer with respect to its shape can be written similarly as previously defined for continuous representations^32,33^ as

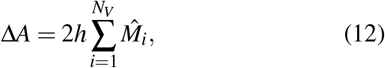

where the sum runs over all vertices and 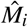 is the mean curvature associated with each vertex, respectively, and *h* is the thickness of the neutral surfaces. When choosing *κ*_Δ*A*_ → ∞ we recover the bilayer couple model where Δ*A* acts as a constraint^33^. Numerically we convert this by choosing *κ*_Δ*A*_ to be on the order of 10^5^ - 10^6^*k*_B_*T*, where *k*_B_ is Boltzmann’s constant and *T* is the temperature^23^.

The overall tether-energy is given by a coupling constant *κ*_*T*_ times the sum over the tether pair-potential *T* (*r*_*ij*_) over all edge lengths *r*_*ij*_,

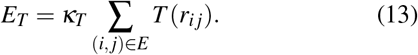

This energy serves to constrain the edge length *r*_*ij*_ and to guarantee the efficient and accurate simulation of fluid triangulated surfaces. Previous work^17,23^ used discrete flat bottom potentials; however, these are not amenable to smooth time integration. In order to use the tether potential in hybrid MC simulations a continuous representation is required^18^. To be able to use large integration time steps Δ*t*, it is crucial to avoid diverging branches that are present in previous formulations. Therefore, we use a continuous tether potential of the functional form

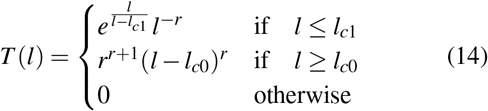

with the *l*_*c*1_ and *l*_*c*0_ being the lower- and upper onset of penalization and the slope *r* ∈ ℕ_1_ of the potential well.

The overall repulsion energy is given by

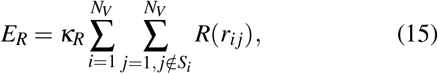

where *S*_*i*_ is a set of excluded vertices for vertex *i*, e.g., the set of directly connected neighbors. The 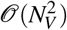 complexity involved in the evaluation of this potential is in practice reduced by the use of efficient neighbor tracking techniques^34^. Mimicking the repulsive electrostatic membrane-membrane interactions, mesh intersection is prevented by the introduction of a penalty that applies a repulsive force on pairs of non-bonded vertices that are below a certain threshold distance *l*_*c*1_. Such a penalty is implemented by the repulsive branch of the tether penalty, Eq. (14). It is given by

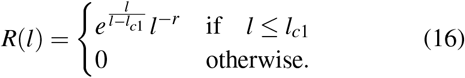

The functional form of the membrane-membrane interaction could be adapted to model various properties of membrane adhesion and repulsion.

## III. ALGORITHMS AND IMPLEMENTATION FOR SIMULATION AND ENERGY MINIMIZATION

In the following we discuss how to efficiently sample the configurational space of the DTS using the hybrid MC approach. First, we introduce the general sampling strategies and discuss how they are implemented. Next, we highlight the issues and solutions for efficient parallelization and evaluate their performance. Finally, we discuss important features to efficiently equilibrate and minimize structures in practice. The different algorithmic components introduced below are shown in the flow-chart in Fig. 1. We apply these tools in Sec. IV.

**FIG. 1.**
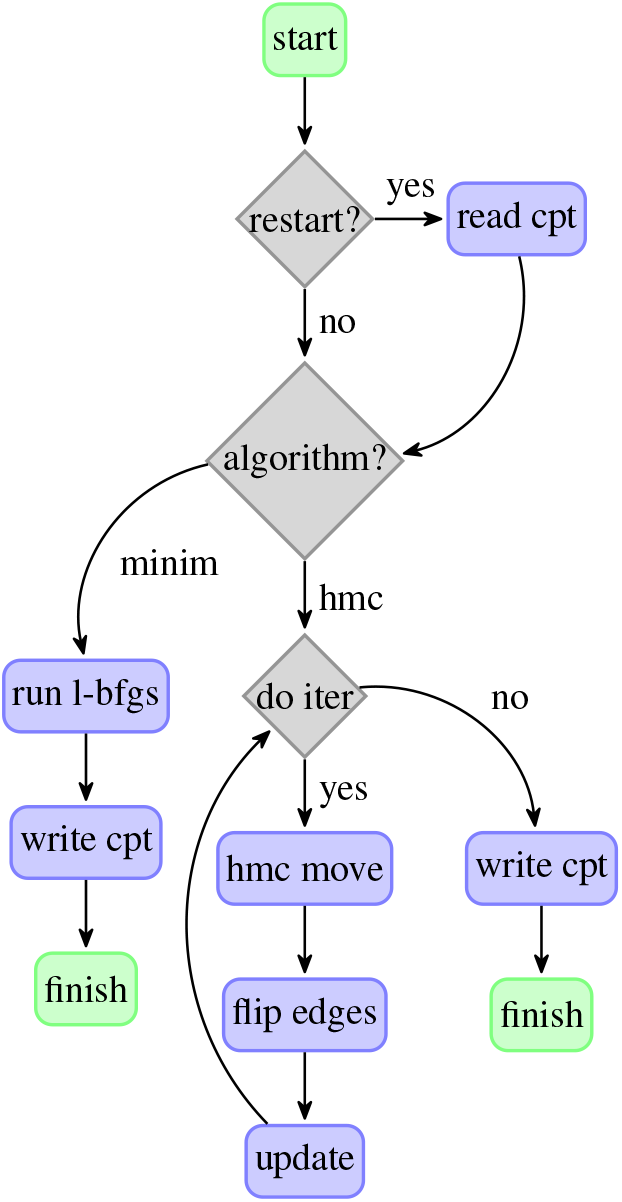
Algorithmic flow-chart of the TriMem software. ‘run l-bfgs’ refers to section III F, ‘hmc move’ and ‘flip edges’ are introduced in Algorithm 1 and Algorithm 2, respectively. ‘update’ summarizes the update of the reference parameters *η*, see Sec. III E, and the temperature *T*, see Sec. III G.

### A. General Sampling Strategy

The statistical properties associated with the Hamiltonian (2) follow the canonical distribution function given by

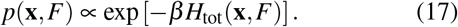

where we use *β* = 1 for the inverse temperature using reduced units. We use a Markov chain MC procedure to sample configurations (**x**, *F*). To account for the compound nature of the state of the triangulation, given by vertex position as well as vertex connectivity, we generate new samples using the Metropolis algorithm with two alternating, well established MC moves: (i) global vertex displacements and (ii) edge flips. After each step the resulting configuration is accepted or rejected according to the Metropolis criterion^15^ by evaluating the differences in the Hamiltonian *H*_tot_ as a function of **x** and *F*. Edge flips ensure membrane fluidity.

Vertex displacements are generated by a Hybrid Monte Carlo scheme (HMC)^35^. HMC draws on the lifting of the vertex coordinates **x** onto an artificial phase-space **x** ↦ (**x, p**), where the components of the vector **p** are the momenta for all vertices. On this phase-space, we impose a symplectic structure via the lifted Hamiltonian

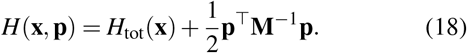

The transpose of a vector is indicated by superscript ‘^T^’ and **M**^−1^ is the inverse of the diagonal mass matrix **M**. We exploit HMC to generate state transitions (**x**_*n*_, **p**_*n*_) → (**x**_*n*+1_, **p**_*n*+1_) in the high-dimensional phase-space with high efficiency. Transitions generated in this way can then trivially be un-lifted (**x**_*n*+1_, **p**_*n*+1_) → **x**_*n*+1_ to give a new sample **x**_*n*+1_. In practice, we use symplectic time-integration schemes such as the leap-frog/Verlet integration method^34^. We show the general concept of a HMC step from *n* → *n* + 1 in Algorithm 1. Verlet-type integration accurately conserves *H*(**x, p**). As a result, the global vertex moves of HMC are accepted with a probability close to one.

Conceptually, a HMC step consists of a short molecular dynamics (MD) simulation of the vertices, with velocities drawn from a Maxwell-Boltzmann distribution and forces given by the negative gradient of *H*_tot_ with respect to the vertex positions (see appendix A). Subsequently, the lifted Hamiltonian is subject to the Metropolis criterion. The time integration parameter Δ*t* and *L* can be used to tune the efficiency of the HMC algorithm. The time step Δ*t* influences the acceptance probability within the Metropolis criterion. The number of steps *L* affects the mixing of the generated Markov chain. Tuning of these parameters is crucial for an efficient sampling scheme. The mass matrix **M** is another free parameter that can be tuned to optimize the sampling performance. In this work we use a single mass *m*, **M** = *m***I**, and set *m* = 1 as a default. This setting has proven efficient in practice. For sampling according to Eq. (17), the application of the HMC framework leads to an enormous gain in efficiency compared to a sequential single-vertex-move dynamic. In particular in a high dimensional regime, i.e., large number of vertices *N*_*V*_, HMC is essential for efficient sampling.

#### Algorithm 1

One step of the HMC algorithm. We randomly draw new momenta from a Maxwell-Boltzmann distribution, perform a number of integration steps, and accept or reject the resulting configuration.

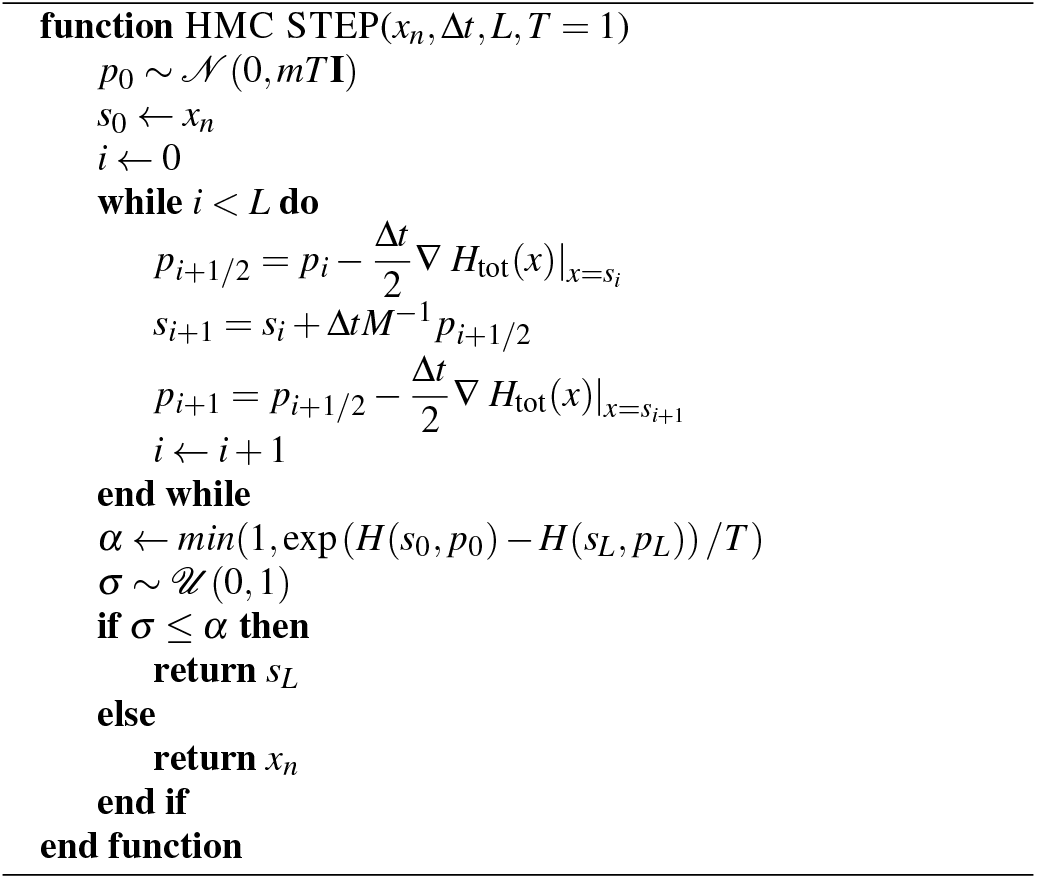

Edge flips ensure the 2D-fluidity of the membrane and alleviate a possible bias due to a fixed vertex connectivity. In an edge flip, the shared edge (*i, j*) of two adjacent triangles (*i, j, k*) and (*i, l, j*) is changed to (*k, l*), resulting in modified triangles (*i, l, k*) and (*j, k, l*) (Fig. 2).

**FIG. 2.**
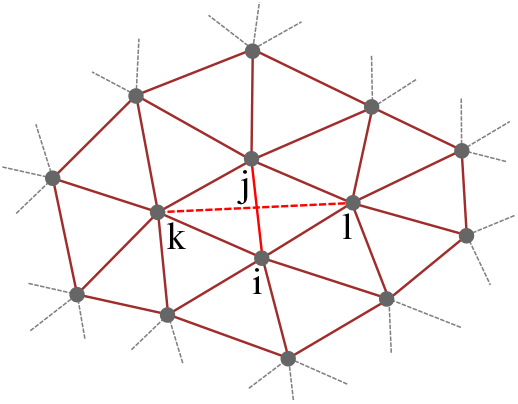
Extent of an exemplary flip patch. Shown is a patch of vertices involved in a flip which is locked during a the flip move. In a flip move the edge (*i, j*) (light red, solid) is flipped to (*k, l*) (light red, dashed). All edges involved in the computation of the averaged vertex properties (dark red) for the vertices *i, j, k* and *l* are included in the flip patch and must be locked during the flip operation. Note, for comparison, that this patch is significantly larger than the patch required for a flip subject to the Delaunay criterion^41^ where the first shell neighbors are not required.

One step of edge flips consists of a sweep over a predefined fraction *γ* ∈ [0, 1] (in the following also referred to a target flip rate) of the set of edges *E*. It is conceptually depicted in Algorithm 2.

### B. Implementation

The vast part of the computational workload of the algorithms is the evaluation of the Hamiltonian and its gradient (see also Appendix A). This evaluation is needed from within the vertex moves as well as in the edge-flip move routine. At the core of the energy computation is the evaluation of the vertex-averaged quantities in Eqs. (4) and (5). Efficient evaluation of the vertex-connectivity is thus essential for overall performance. To this end, we utilize the concept of the half-edge data structure. This data structure for polygonal meshes efficiently tracks incidence information of edges, vertices and faces^36,37^. In particular, we make use of the OpenMesh library, which provides a generic implementation of the half-edge data structure in C++^38^ that can handle various mesh sizes and geometries as input. To fully leverage the OpenMesh interface, we implement the evaluation of the Hamiltonian and its gradient in C++ and provide bindings to Python using the pybind11 Python package^39^. This approach combines the efficiency of a compiled programming language with the convenience of the well-established Python ecosystem for clear and extensible algorithm development. Using C++ for the computationally intensive evaluations further allows us to exploit shared-memory parallelism and the speedup offered by modern multi-core architectures. To achieve this, we use OpenMP^40^ to parallelize the loops that involve scans or reductions over all vertices in the triangulation, such as Eqs. (6)- (10) and (12) and the respective gradient evaluations (see Sec. III D). While parallelization of both the vertex moves and components of the integration of short trajectories is straight-forward, flip moves are more complex and require a more detailed discussion.

#### Algorithm 2

One sweep of edge flips. The input is the set of triangles *F*, the set of edges *E*, and the fraction *γ* of flips to attempt. It makes use of the Hamiltonian *H*(*F*) as a function of the mesh triangles, and the routines shuffle_edges(*E*) to shuffle the list of edges and flip_egde(*e*), which takes an edge, flips it, and alters the mesh connectivity. The routine called update_hamiltonian(*h, F, e*) incrementally updates the total energy for the changed flip patch (see Fig. 2).

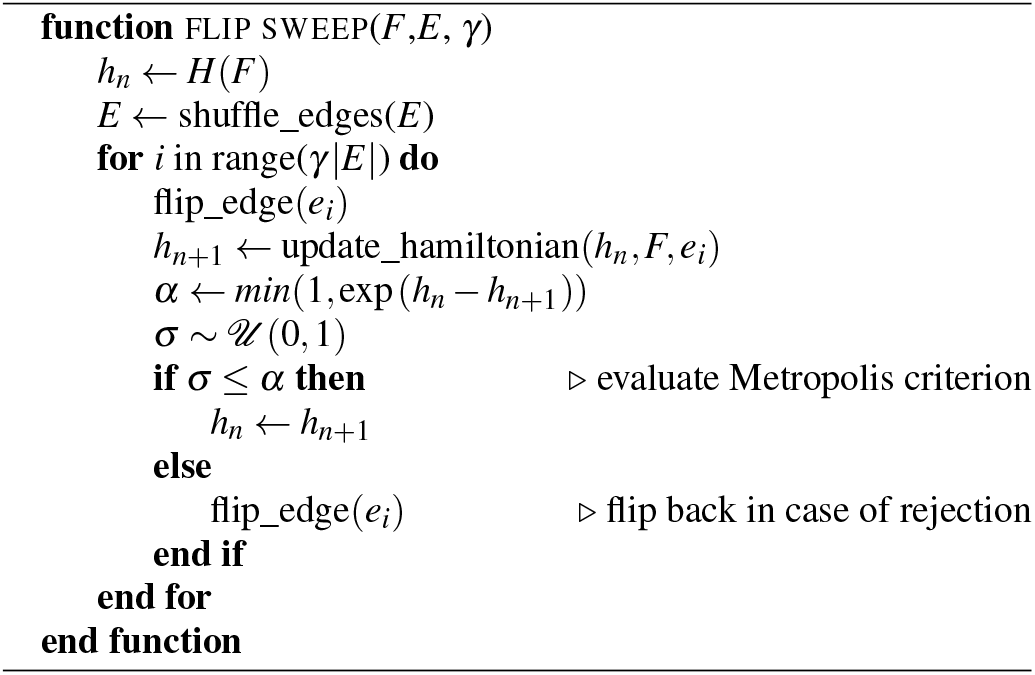

### C. A strategy for the parallel evaluation of edge flips

To increase the computational efficiency of flip moves, we pre-select a set of possible flips and evaluate in parallel independent changes of geometric membrane properties and of the energy. We then evaluate sequentially the corresponding Monte Carlo flip moves, including the associated energy changes from the aforementioned pre-calculated quantities. We designed this Algorithm 3 to alleviate two of the main efficiency bottle necks of Algorithm 2: (i) flip execution, i.e., the technical realization of the change in vertex connectivity that is provided by the underlying data structure; and (ii) energy evaluation, i.e., the evaluation of the change in the Hamiltonian due to the change in connectivity. Both components are crucially influenced by the evaluation of the vertex connectivity.

Due to the edge-based connectivity information implemented by the half-edge data structure, flip execution provided by OpenMesh is realized efficiently by simply swapping edge connectivity^38^. Since the individual edge flip can technically be performed efficiently, the main computational workload results from the energy evaluation of the flip-patch associated to an edge flip. A straightforward exploitation of the speed-up provided by the evaluation of the change of the Hamiltonian for several edge flips in parallel is, however, still complicated by two intermingled issues.

First, individual edge flips tend to have very low acceptance probabilities due to the rather strict penalty on neighborhood distances that is imposed by the tether potential (Eq. (14)). The evaluation of the Metropolis criterion for a flip of several edges at once thus suffers from an exponential decay of the acceptance probability due the multiplicative nature of the acceptance probabilities of single flips. Consequently, the acceptance of edge flips must be evaluated sequentially to establish a reasonable flip rate *ε*, which is defined as

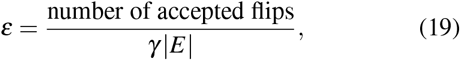

where |*E*| indicates the cardinality of the set of edges *E*. This sequential evaluation therefore represents an inherent serial component of Algorithm 2 that limits the potential speed-up by construction.

Second, selected edges have to be independent such that we can evaluate the respective changes in the Hamiltonian in parallel. Afterwards, we sequentially evaluate the respective acceptance probabilities. The independence of edges means that they cannot be part of the same flip patch. A flip patch is the set of edges affected by a flip (see Fig. 2). This condition imposes a constraint on the set of edges for which the change in the Hamiltonian can be evaluated in parallel. In order to avoid the NP-hard problem of a deterministic pre-computation of sets of non-interfering edges, we propose a randomized approach that is outlined in the remainder of this section.

Inspired by the batch-parallel evaluation of mesh properties of Shang *et al*.^42^, we adopt a batch-parallel version of the flip-sweep Algorithm 3. Edges are first linearly assigned to participating threads by chunking the vector of edges as managed by the underlying data structure. During the sweep-iteration, edges are then selected at random by each thread from its respective chunk. To ensure independence of such a parallel batch of edges, each edge is attempted to be locked together with the associated flip-patch for the use by other threads. For technical reasons, we impose this strict condition of non-overlapping flip-patches within a batch of edges even though non-inclusion of a flipped edge in another patch would already suffice. The implementation draws upon a scoped data locking mechanism that allows for an efficient detection of patch clashes. Upon success of locking, a thread can independently evaluate the patch-local contributions to the geometric properties *M* (Eq. (6)), *A* (Eq. (9)) and *V* (Eq. (10)), as well as the contributions to the bending energy (Eq. (3)), the tether-penalty (Eq. (14)) and repulsion-potential (Eq. (16)). The computationally intense evaluations are thus parallelized over a batch of edges with batch-size equal to the number of threads used in parallel. Subsequently, the change in the Hamiltonian and the acceptance probability for each edge in the batch are evaluated sequentially, thus preserving ergodicity of the resulting Markov chain. Although this inherent serial component reduces the theoretically achievable speedup as mentioned above, it has shown to be satisfactory in practice while maintaining a flip rate *ε* close to the serial version (see section III D for detailed results).

### D. Parallel performance

We analyze the parallel performance of the algorithms and methods introduced in Sec. III on a unit sphere with different degrees of mesh discretization as a test geometry. We use the icosahedron recursion technique from the meshzoo Python package^43^ to create high quality meshes with *N*_*V*_ ∈ [624, 2562, 10242, 40962, 163842, 655362] vertices. The results referred to in this section are produced on a dual socket node with Intel® Xeon® Platinum 8280 CPUs with 56 cores in total.

Figure 3 shows results for the parallel scaling of the different algorithmic components with the number of threads for different numbers of vertices: energy evaluation (Eq. (2)), its gradient/force, flip-sweep according to Algorithm 3 and a full step of the MC-procedure outlined above, comprised of 1 HMC-step (Algorithm 1 with *L* = 100 and Δ*t* = 1.0×10^−4^; note that the results in Fig. 3 and Fig. 4 are invariant with respect to the exact value of the time step Δ*t*) plus 1 flip-sweep. To improve consistency, the measurements for energy, gradient and flip-sweep comprise 10 evaluations of the respective components due to their short run-time.

**FIG. 3.**
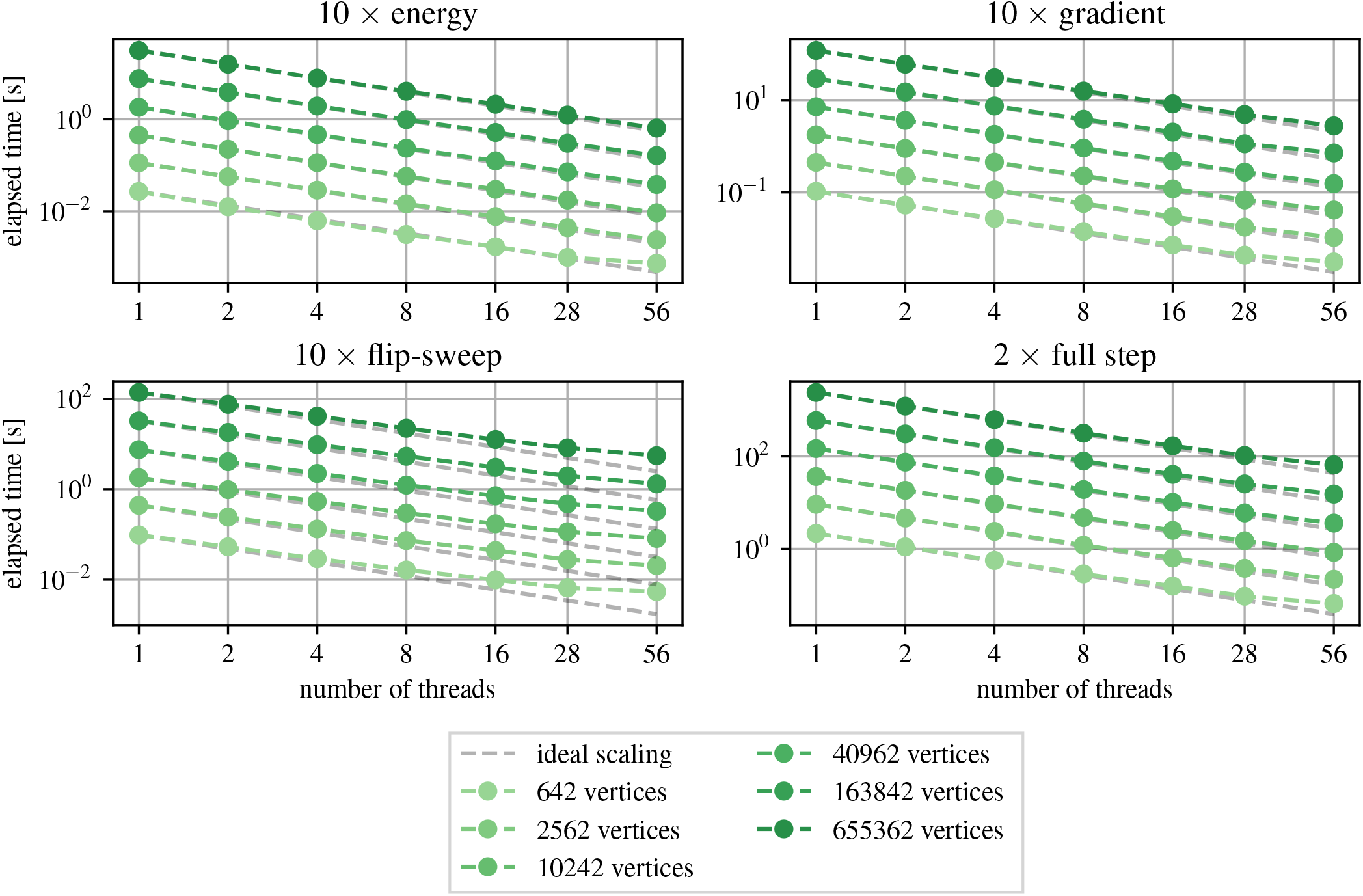
Strong scaling of different algorithmic components with the number of threads for different problem sizes *N*_*V*_ (shades of green). The components are the evaluation of the energy, Eq. (2), its associated gradient/force, a flip-sweep according to Algorithm 3, and a full step of the MC-procedure outlined in Sec. III with 1 HMC-step (with *L* = 100) and 1 flip sweep. For the components energy, gradient, and flip-sweep, measurements consist of 10 evaluations each. The measurement for the component full-step consists of 2 steps.

**FIG. 4.**
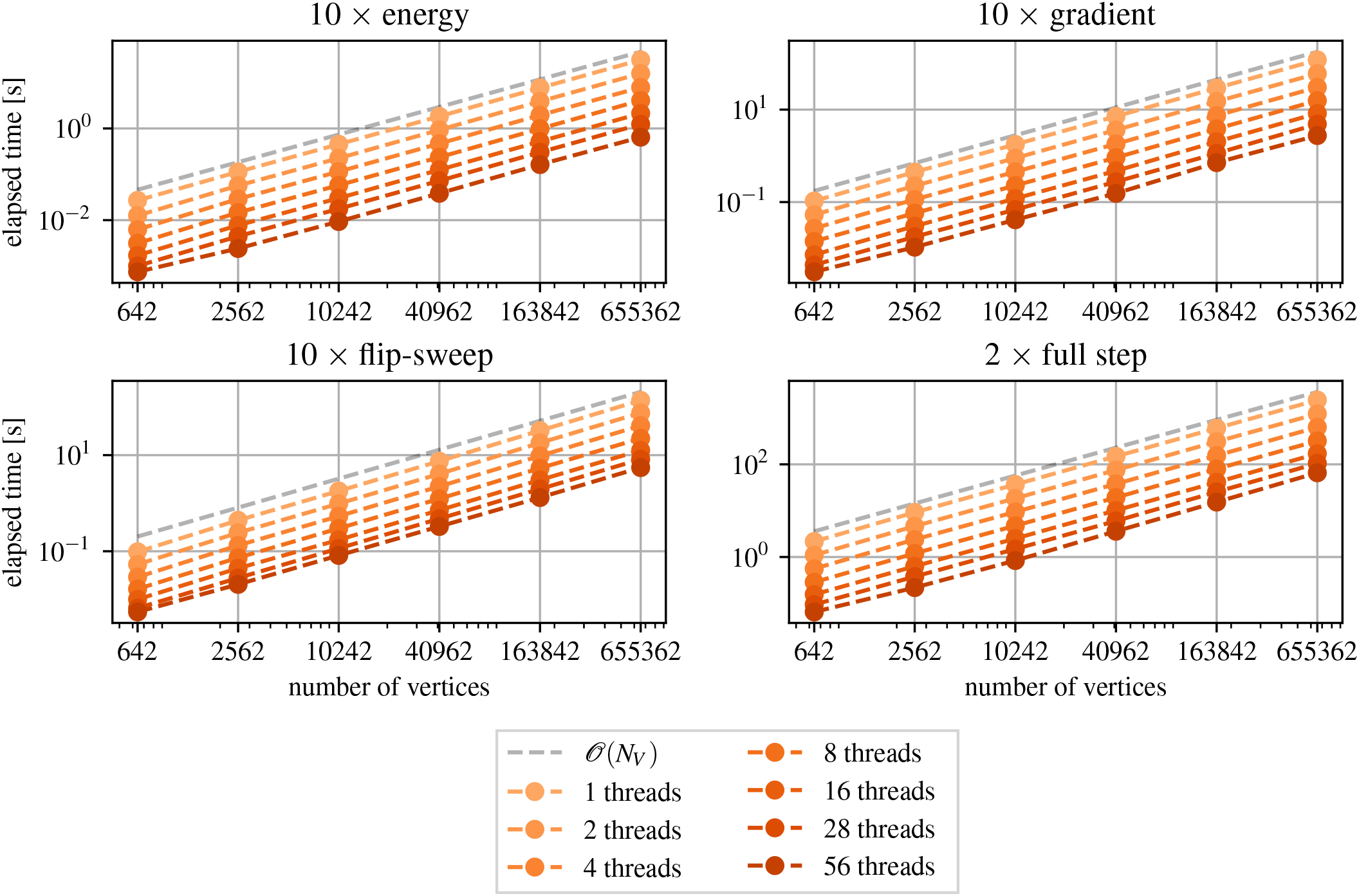
Time complexity of different algorithmic components with the number of vertices *N*_*V*_ for different numbers of threads (shades of orange). The components are according to Fig. 3. For the components energy, gradient and flip-sweep, measurements consist of 10 evaluations each. The measurement for the component full-step consists of 2 steps. Color intensity indicates higher thread numbers. The slope of the grey dashed line indicates 𝒪(*N*_*V*_) linear time.

The high arithmetic intensity involved in the energy and gradient evaluations leads to efficient use of the available resources, which is reflected in good parallel speedup on the compute node. The flip-move sweep and MC-step evaluations also benefit from an increase in parallel execution performance, but a saturation trend is visible. This trend is a result of both of these components containing a significant amount of low arithmetic intensity workload. Thus, they are more exposed to the bound of the memory bandwidth than the energy and gradient evaluation. In addition, the flip-sweep still has a serial portion that will inherently limit the achievable speedup. However, in the experiments presented here, this does not manifest itself as the critical component controlling the effective speedup. The scaling of the full MC-step is only minimally affected by the flip-sweep due to the small contribution of the flips-sweep to a full MC-step. Instead it is governed by the amount of low arithmetic intensity workload in the HMC-step (Algorithm 1).

#### Algorithm 3

Parallel version of algorithm 2. The directives **# BARRIER** and **# CRITICAL** refer to OpenMP directives used in shared memory parallelism^40^. **# BARRIER** indicates a barrier where threads have to wait for each other. **# CRITICAL** defines a region that can only be executed by one thread at a time.

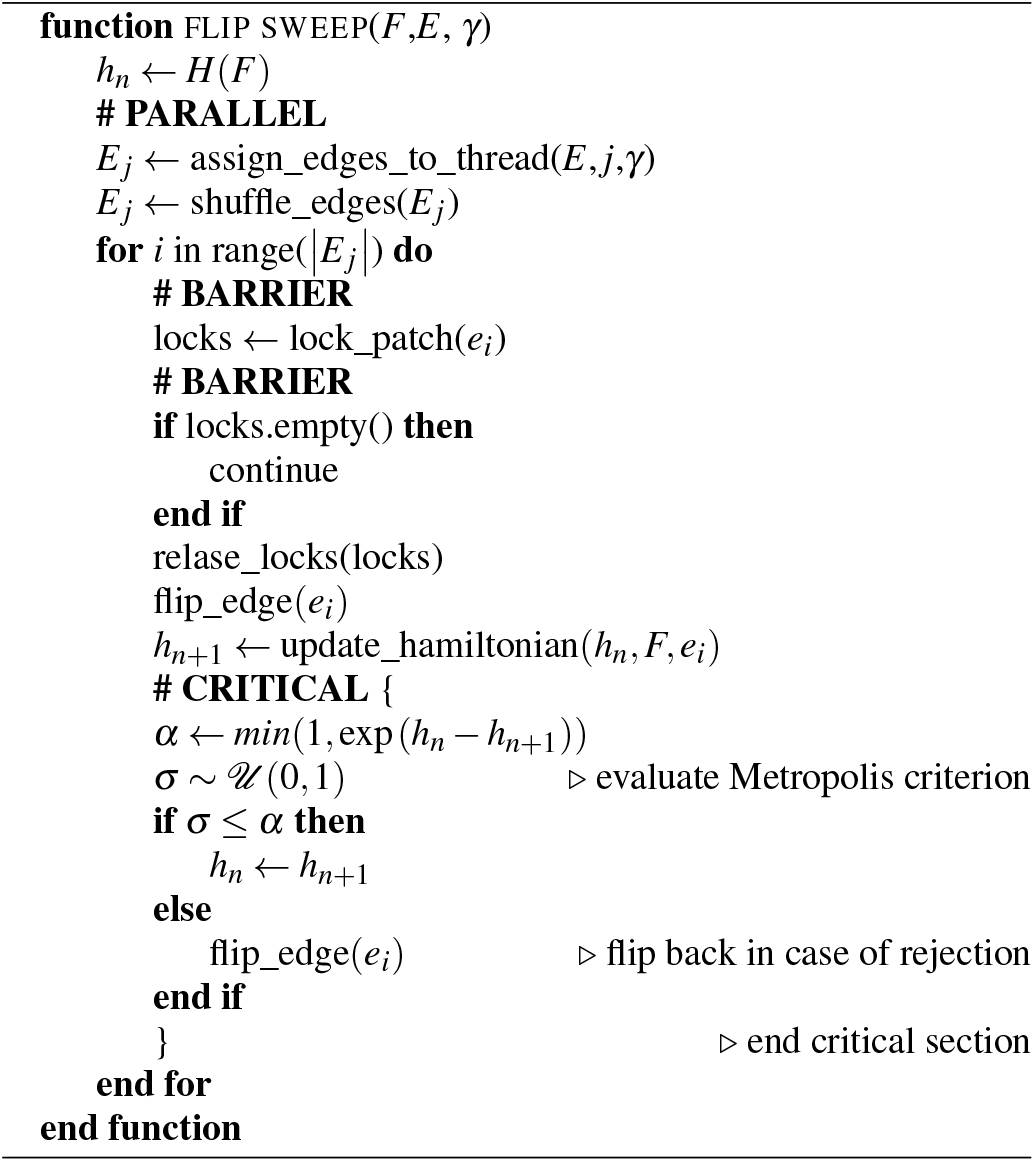

Importantly, with our parallelized scheme, the computation of a mesh with 40962 vertices is comparable to a single-core run with a small system with 642 vertices. Without an explicit limit on mesh size in openmesh, it is possible in principle to simulate meshes at least up to a size of 10^6^ − 10^7^ vertices (Fig. 3), as required for cellular scale membranes. The global vertex moves in HMC ensures efficient sampling of the high dimensional space even for large mesh sizes.

The effective time complexity of the algorithmic components is shown in Fig. 4. The distance calculations necessary for the computation of the repulsion penalty given by Eq. 16 are effectively reduced to 𝒪(*N*_*V*_) by the neighbor list data structures. This complexity on the level of the energy and gradient evaluation is directly passed on to the evaluation of the full MC-step.

The influence of the parallel implementation (algorithm 3) on the flip rate *ε* of the flip-sweep is shown in Fig. 5. The measurements are carried out for a mesh with *N*_*V*_ = 10242 vertices and a target flip rate *γ* = 10%. We found that the mean efficiency of *ε* ≈ 0.17% observed for the serial implementation reduces to ≈ 0.15% when using a full node with 56 cores. This slight decay is due to our randomized approach, in which the number of clashes in the locking of the necessary flip patches increases with the number of parallel threads used. These clashes lead to threads skipping a flip-batch and can consequently result in a reduced number of edge flips in total. Due to the linear distribution of edges to threads, the spatial order of edges as reflected in the data structure in memory can have an influence on the probability of clashes. As applied by^42^, a spatial pre-ordering of the edges (e.g., by space filling curves) might thus improve the efficiency for high thread counts. Since the spatial ordering of vertices resulting from the icosahedron reconstruction used here is already rather good, preliminary tests have not shown to improve the presented results. Nevertheless, for general meshes the influence of spatial ordering is considered to have significant influence. Therefore, this topic might be followed up in future developments. In any case, the obtained effective flip rates have proven sufficient to provide the desired effect on the mesh mutability that is necessary to achieve membrane fluidity and accurate results, cf. section IV.

**FIG. 5.**
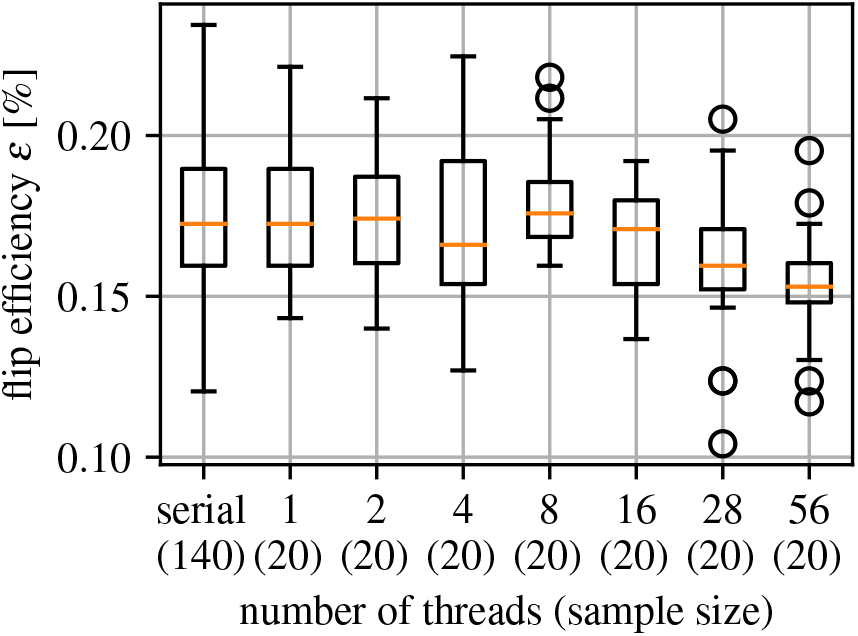
Flip rate *ε* given by Eq. (19) of the parallel flip implementation given the number of threads. The target rate is *γ* = 10% and the number of vertices is *N*_*V*_ = 10242. For comparison, we show the rate for the serial version (leftmost data). The cardinality of the sample size is given in parentheses for every box. The flip efficiency decreases slightly with increased number of threads due to the locking of flip patches from different threads.

### E. Parameter Continuation for Stiff Restraints on Membrane Shape

The accurate representation of the constraints regarding volume and surface area requires large values for the factors *κ*_*V*_, *κ*_*A*_ and 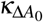 in the respective penalty formulations given by Eqs. (7), (8), and (12). The conditioning of the Hamiltonian in Eq. (2) is determined by these penalty terms. That is, slight deviations of the initial configuration from the desired target configuration in terms of the area *A* and the volume *V* can have a large impact on the numerical stability of the Hamiltonian and the sampling performance. To mitigate such a performance degradation, TriMem offers the possibility to use a technique from the concept of parameter continuation^44^. To this end, we introduce the parameterized Hamiltonian *H*_*p*_(**x**, *F*; *η*), with the explicit definition of parameters *η* := (*V*_0_, *A*_0_, Δ*A*_0_). This enables a smooth blending of the Hamiltonian *H*_*p*_(**x**, *F*; *η*_0_) → *H*_*p*_(**x**, *F*; *η*_1_) from some initial parameters *η*_0_ to the target parameters *η*_1_ via the linear interpolation

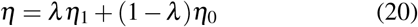

by varying an interpolation parameter *λ*∈ [0, 1] smoothly from 0 → 1. By doing so, the sampling efficiency remains higher, even in situations in which the initial configuration is not consistent with the specific constraints in place. Parameter continuation can also help to overcome energy barriers that might appear in an immediate instantiation of the target Hamiltonian. More generally, the parameterized Hamiltonian allows us to integrate systematic approaches to branch tracking^44^ in future work.

Currently, we interpolate the whole set of parameters *η* simultaneously by defining and implementing a single interpolation parameter *λ*. The efficiency of this scheme could be improved by using a vector of interpolation parameters such that individual components of the Hamiltonian are transformed independently.

### F. Gradient-based energy minimization

If the initial configuration is not consistent with the imposed constraints, we can improve the sampling efficiency by initial energy minimization. Such a minimization can be interpreted as the one-time application of a preconditioning and will bring the initial configuration closer to the equilibrium configuration, determined by Eq.(17). Energy minimization also avoids the necessity of running long simulations with parameter interpolation in the beginning. Since this is only an initial energy minimization prior to a HMC simulation, no particular requirements must be imposed on the convergence to the global optimum. By ignoring edge flips during minimization, we can use well established and efficient methods for function optimization such as the L-BFGS method^45^. In TriMem, simulations can be run in minimization or HMC mode (see Fig. 1).

### G. Simulated Annealing

Gradient-based Energy minimization of the vertex positions must be followed by a global minimization strategy accounting for both vertex positions as well as mesh connectivity. We apply a simulated annealing procedure for the exploration of the domain of Eq. (17) that is capable of finding global minima/maxima. Following Ref.^46^, we implement this method by simply modifying the temperature argument of the HMC step in Algorithm 1 according to a cooling schedule. Specifically, we apply an exponential cooling scheme *T*_*n*+1_ = max[exp(−*λ*)*T*_*n*_, *T*_min_], with the cooling factor *λ* ∈ ℝ^+^ controlling the rate of cooling.

In TriMem, cooling can be initiated during HMC simulations directly in the beginning or after a longer equilibration period prior to the cooling. This enables sufficient sampling of the configurational space before settling into the global (or deep local) energy minimum.

## IV. VALIDATION METHODS

We tested the robustness of the TriMem software and compared it to analytical and numerical calculations from the literature. We reproduced several well established aspects of the phase diagram for closed vesicles with *c*_0_ = 0 and with respect to varying volume or area difference. In all simulations, we used a bending rigidity of *κ*_*B*_ = 30*k*_B_*T*, which is a typical value for biological membranes. The coupling constants of the volume, *κ*_*V*_, the area, *κ*_*A*_, and the area difference, *κ*_Δ*A*_, were chosen several orders of magnitude larger with *κ*_*V*_ = *κ*_*A*_ = *κ*_Δ*A*_ = 1 × 10^6^*k*_B_*T* to impose a strong restraint. All simulations were performed using a mesh size of *N*_*V*_ = 1962 or 642 vertices starting from a sphere. The initial shapes were generated with the meshzoo library^43^. Our results are not affected by mesh size.

A practical instruction to setup and run such simulations with TriMem is available via Github (https://github.com/bio-phys/trimem) on the documentation webpage.

### 1. Volume Phase Diagram

The individual configurations of the branches of the volume phase diagram (Fig. 6) were generated as follows. The initial shapes for each branch were generated using the minimization procedure to initialize the system. Prolate simulations were started at the reduced volume *v* = 3*V/*(4*πR*^3^) = 1.0 (Fig 6; blue triangles), oblate simulations at volume *v* = 0.65 (orange circles) and stomatocyte simulations at *v* = 0.3 (green squares). These initial shapes and volumes on the respective branches were achieved by using the preconditioning procedure (see Section III F). Then, using these initial shapes the reference reduced volume *v* was lowered or increased by 0.025 instantaneously in each step. In each step the previous final structure was used as initial configuration for the subsequent step. By doing so the respective branches could be mapped out by exploiting hysteresis. In all cases the simulations could, in principle, switch their respective branches on the phase diagram and otherwise no special restraint were applied. Each simulation was run for 5 × 10^5^ steps using a temperature *T* = 1*k*_*b*_*T* and in an additional 6 × 10^5^ steps the temperature was reduced from 1 to 0 following section III G. The rescaled bending energy of the lowest energy configuration was plotted for each value. In all HMC simulations an integration step size of 7^−5^ was used. The trajectories in each HMC step were 100 steps long. The flip ratio was set to *γ* = 0.1 of the vertex move steps. Previous tests showed this flipping ratio was sufficient for fast vertex diffusion.

**FIG. 6.**
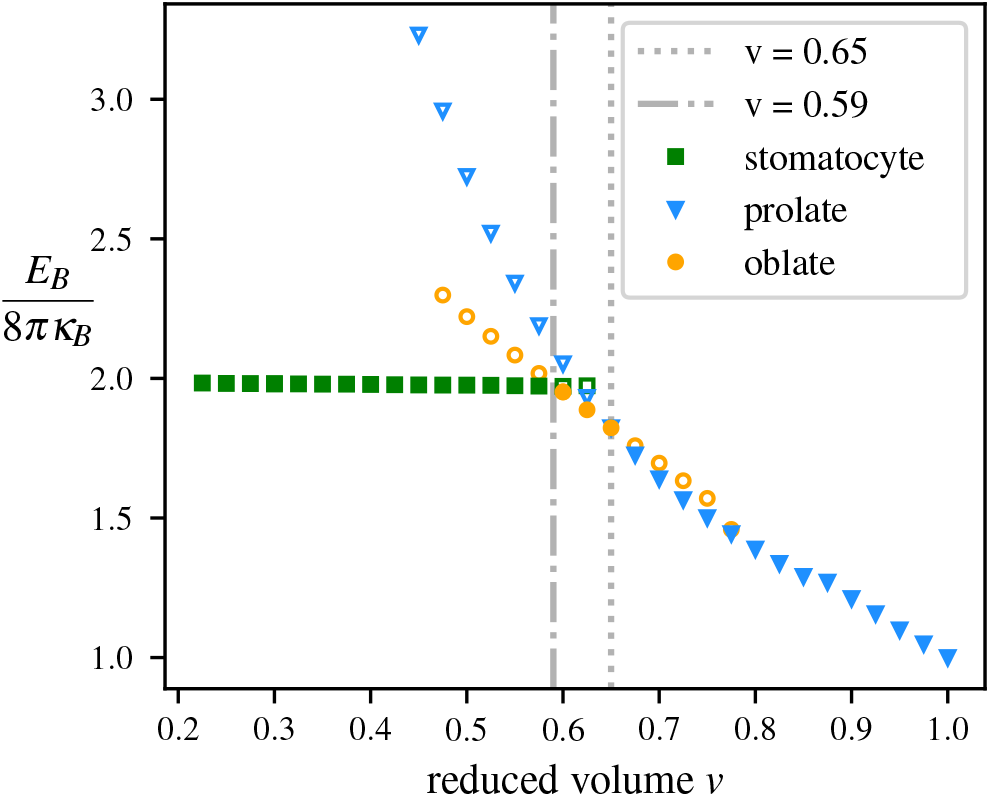
Phase diagram for a vesicle with *N*_*v*_ = 1962 vertices of spherical topology in the plane of the reduced volume *v* and the bending energy *E*_*B*_. Shown are minimum energies obtained for different membrane shapes as a function of *v*. Filled symbols indicate the respective lowest-energy shapes. Open symbols indicate metastable shapes at a given reduced volume. All states with prolate shapes are shown as blue triangles. Oblate shapes are shown as orange circles. Stomatocytes are shown as green rectangles. The dot-dashed and dashed lines correspond to *v* = 0.59 and 0.56, respectively, and mark transitions between stable branches. Exemplary energy-minimized shapes for all corresponding branches are shown in Fig. S1.

### 2. Area-Difference Phase Diagram

The area-difference phase diagram is another commonly used benchmark to test the robustness of numerical membrane bending simulation codes^23,28,47^. The simulations were performed as follows. The initial shape for the HMC run was generated in a two-step minimization procedure. First, the volume was reduced to the respective target reduced volumes of *v* = {0.5, 0.55, 0.6}, and second, the reduced area difference Δ*a* = Δ*A/*(4*πR*) was adjusted to the respective target values between 1 and 1.8, by using the L-BFGS minimizer for gradient based energy minimization for both parameters successively. Then a HMC simulation was run for 1.5 × 10^7^ steps. After 1.4 × 10^7^ steps 1 × 10^6^ steps of cooling were performed where the temperature was reduced from 1 to 0 according to section III G. The bending energy and shape at the final step of each simulation were extracted and normalized to the reduced energy by 8*πκ*_*B*_ to construct the phase diagram. Otherwise the same parameters were used for the HMC simulations as described above in section IV 1.

## V. RESULTS AND DISCUSSION

We extensively validated our software for varying mesh sizes and exhaustively tested all features by reproducing known phase diagrams of vesicle shape^47,48^. First, we ran a range of simulations to produce the volume phase diagram for a vesicle shown in Fig. 6. In the following, we consistently use reduced volumes *v* as an order parameter. Starting with a sphere (*v* = 1), we explored the range between *v* = 1 and 0.2 to locate minimal shapes found as described in section IV. The *y*-axis shows the bending energy *E*_*B*_ and was re-scaled with 8*πκ*_*B*_, the bending energy of a sphere, as in previous work^23,47^.

Here, we explored the stability of the three well established branches in the vesicle volume phase diagram; the prolate, the oblate and the stomatocyte^17,28^. The transitions between the three branches are not smooth, i.e., the first derivative of the energies with respect to the reduced volume are discontinuous. Stable and metastable states representing the various shapes are separated by energy barriers.

The prolate branch (Fig. 6, blue triangles) gives the global energy minima for *v* ≿ 0.65. As the reduced volume is lowered, the vesicle changes its shape from the sphere at *v* = 1 to a prolate shapes and then to extended tubular structures. Low reduced volumes result in long metastable tubes with narrow diameter and a far higher bending energy than the corresponding oblate or stomatocyte shapes with the same reduced volume, which constitutes the global minimum for *v* ≾ 0.65. Below this reduced volume the branch is metastable as previously outlined in^23,28^. All prolate shapes and tubes below *v* = 0.65 are only found when starting from an initial prolate configuration and mapping out the branch by hysteresis, indicating that they are local minima and metastable.

The oblate branch is the global minimum between 0.59 ≾ *v* ≾ 0.65 (Fig. 6 orange spheres). Simulations starting from the sphere instantaneously reducing the volume and using the L-BFGS minimizer to a target value *v* < 0.65 always converged to an oblate shape. This result is in agreement with observations of previous work^17,28^. Above *v* = 0.725, the oblate is always unstable and the local energy minimum for the branch vanishes, and all initial shapes (sphere, prolate, oblate) converged to the prolate shape, which is consistent with previous theoretical work^49^. Oblates in the range of 0.65 < *v* ≲ 0.725 can only be seen when starting with an oblate configuration in the globally stable range and tracing out the branch by hysteresis. By doing so the structures remaining in the metastable minimum. Similarly, the oblate shape is metastable at a reduced volume of *v* ≲ 0.59.

For a reduced volume of *v* ≲ 0.59 the stomatocyte is the global minimum, which is also achieved from the preconditioning step with the L-BFGS minimizer from a sphere using a stiff restraint on the target volume (Fig 6 green squares). For *v* > 0.59, we find that the stomatocyte shapes are metastable with respect to oblates. The energy of the stomatocytes are nearly independent of the reduced volume with *E*_*B*_ ≈ 16*πκ*_*B*_. Overall, all energies for the globally energy-minimized shapes are in good agreement with previous work following similar approaches^17,28^.

To test if the area-difference is correctly evaluated, we also produced the phase diagram for varying area differences at multiple fixed reduced reduced volumes. Fig. 7 shows the rescaled bending energy as a function of the area difference Δ*a*_0_. Simulations were run for area-difference increments of 0.025 and the energy-minimized structures after cooling to *T* = 0 were plotted (see IV).

**FIG. 7.**
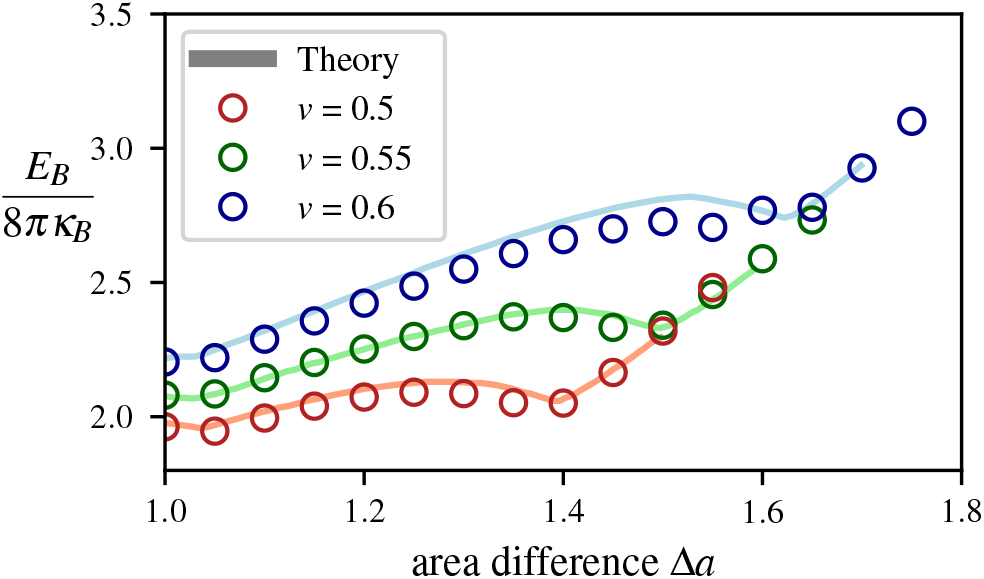
Energy as function of reduced area difference Δ*a* at constant volume *v* for a mesh with *N*_*v*_ = 642. The simulation results are shown as empty circles for a constant volume *v* = 0.5 (blue), 0.55 (green), and 0.6 (red), respectively. For reference, the light lines are adapted from earlier calculations^47^ using numerical triangulation^50^. Exemplary energy-minimized shapes for varying area difference are shown in Fig. S2

We verified that both the minimized energies and the corresponding shapes are in agreement with previous simulations and theoretical calculations^23,28,47^. Small differences in the bending energies for some shapes in Fig. 7 are likely caused by hysteresis effects around boundaries in shape space, the use of edge flips here, and different resolutions of the triangulations. The oblate configuration and the tube shapes are metastable for *v* = 0.55 and are separated by an energy barrier. The top of the barrier corresponds to the non-axisymmetric paddle shape which separates the prolate and tube branch and converts to a tube with increasing Δ*a*. For all values with Δ*a* < 1.4 we observed structures with a *D*_3*h*_ symmetry. First triangular shaped oblates emerge which transition into three-armed starfish shapes (Fig. S2). We also find that the dumbell/tube structure^47^ is a local minimum for Δ*a*_0_ > 1.4. Tubes are found for all higher values of Δ*a*_0_. We note that extensive descriptions and discussions of these phase diagrams and their symmetries are provided in previous systematic studies^23,28,47^.

Taken together, these simulations validate that TriMem faithfully reproduces data from previous studies and can be used to study a broad range of reshaping processes. All parameters are correctly assessed and our parallel dynamics protocol produces the expected results.

## VI. CONCLUSIONS AND OUTLOOK

We presented TriMem, a parallelized open-source software package for hybrid MC simulations of triangulated meshes representing lipid membranes. TriMem offers a robust, efficient, and parallel implementation of the method that enables users to perform simulations efficiently with large mesh sizes of 10^6^ − 10^7^ vertices, which was not previously possible. Many previously available software packages were limited to a few thousand vertices by the computational cost. We demonstrated how all relevant move types can be efficiently parallelized across multiple CPU cores up to a full two-socket shared memory compute-node (Fig. 3). While parallelization of energy and force routines to generate short MD trajectories for vertex moves is straightforward, we showed that flip parallelization requires a more complex algorithm to produce correct results. We introduced a novel method for flip moves that combines an effective generation of sets of edges with non-interfering flip-patches. The parallel computation of the independent energy changes and subsequent evaluation of the acceptance in series increases the performance. We bypass a major issue resulting from serial energy evaluation required for individual flip moves.

Additionally, the HMC approach allows for more efficient global vertex displacement moves with acceptance ratios of nearly one due to efficient proposals generated by short molecular dynamics trajectories of the mesh and vertices^18^. This leads to fast convergence of sampling and will be particularly useful when simulating large meshes with millions of vertices.

We implemented an efficient neighbor-list algorithm to enable the use of repulsive, and possibly adhesive potentials, between non-neighbor vertices. The resulting code scales linearly, 𝒪(*N*_*V*_), with mesh size, and makes calculations with large meshes possible. This feature proved particularly important to sample highly curved membrane shapes such as stomatocytes, where self-intersection of the mesh could result in faulty representations. It can also be further expanded to model the general physical properties of membrane repulsion due to charges, possible protein-mediated adhesion, and systems with multiple membrane meshes.

The user interface of the software was designed for versatility but with ease-of-use in mind. Some functionality, such as the configuration file, is kept loosely in the style of well established molecular simulation packages such as GROMACS^51^, which is widely used in the biophysics community. The ease-of-use and open-source nature of the code should empower a broad community of biophysicists to quickly set up, run, and evaluate large-scale membrane remodeling processes of interest. The Python front-end should also enable other users to quickly add novel features to the code.

We validated the software by reproducing several phase diagrams and results for vesicle shaping, which have been previously explored in various software packages and foot on theoretical calculations^17,23,28,47,49^. We found in all cases that the validation was in good agreement with existing work, indicating the robustness of our algorithms.

The code was designed for performance with C++ in the backend and for ease-of-use with a Python interface in the frontend. The framework provided by the TriMem package can be extended in the future in a straightforward way to incorporate more complex biological assemblies of membranes. Protein membrane interactions, multiple membrane meshes, and cyctoskeletal attachments will likely be added to the features of TriMem, enabling a broader range of cell-scale simulations. Systems containing multiple membranes in close vicinity are needed to build organelles and cell-scale membrane structures. Additionally, protein vertices and lipid domains can be included to study complex protein-induced membrane remodeling processes^7,19^.

We anticipate that TriMem, with some modifications, will also enable simulations of complex membrane topologies. One possible direction is to include periodic boundary conditions (PBC) when defining the topology through the local connectivity of the graph. An implementation of PBC will enable the preparation of quasi-infinite membrane shapes from tubes (PBC in *z* direction) over flat membranes (PBC in the *x*-*y* plane) to lipidic mesophases (PBC in all directions *x, y, z*). Another possible direction is to simulate multiple disconnected membranes, such as one vesicle enclosing another, or stacks of membranes as they occur in the Golgi^25^. Here, the vertex repulsion can ensure that the different membranes do not interpenetrate. It is also possible to fix the spatial position of certain vertices. In this way, the dynamic membrane can be connected at fixed seams to other shapes, e.g., to form the half-toroidal pores connected with two flat background membranes as seen in nuclear pore complexes^12^.

In the future, we envision that TriMem in combination with modern structural biology techniques such as cryo-ET will shed light on the structure and dynamics of membranes on an organelle and eventually cell-scale level. Cryo-ET will increasingly serve as data source to build accurate computational models for coarse-grained simulations. We envision a combination of multi-scale simulation techniques including TriMem’s parallelized capability of performing large-scale simulations required to gain mechanistic understanding of cell-scale membrane remodeling processes.

## VII. SOFTWARE AVAILABILITY

TriMem is available as free software under a GPL-3 license from (https://github.com/bio-phys/trimem) and can be installed as a Python package using *pip*^52^. As such, it can be used as a Python library with predefined building blocks for the methods and algorithms presented in the previous sections. It also provides a command line interface that encapsulates the algorithmic flow shown in Fig. 1, which is controlled via an input configuration file. For a more detailed description on the usage, we refer the reader to the documentation which can be found via the Github link above. Software contributions from third party developers are welcome as pull requests on Github.

## SUPPLEMENTARY MATERIAL

See the supplementary material for Figs. S1 and S2 showing minimum-energy vesicle shapes.

## ACKNOWLEDGMENTS

This work was supported by the Max Planck Society. G.H. and M.S. acknowledge funding by the Landes-Offensive zur Entwicklung Wissenschaftlich-Ökonomischer Exzellenz LOEWE of the State of Hesse (https://wissenschaft.hessen.de/wissenschaft/landesprogramm-loewe). M.S. acknowledges support by the EMBL Interdisciplinary Postdoc Programme under Marie Curie COFUND actions.

## Appendix A: Gradient of the Hamiltonian

The gradient of the Hamiltonian (2) can be derived in terms of the derivatives of quantities defined on the n-simplexes that define the triangulation *𝒯*. For the derivatives of these quantities with respect to the vertex positions *x*_*i*_, we apply efficient and well-known formulas from discrete geometry^53^. In particular, we need the derivatives of the edge length, the face area, the face volume and the dihedral angle with respect to the involved vertex coordinates.

For the length *r*_*i j*_ =‖*u*‖ = ‖*x* _*j*_ − *x*_*i*_‖ of the directed edge (*x*_*i*_, *x* _*j*_), the gradient is given by

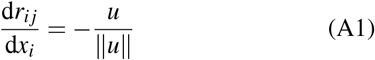

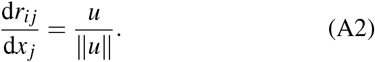

The gradient of the area *A* of an oriented triangle (*i, j, k*) ∈ *F* with normal *n* is given by

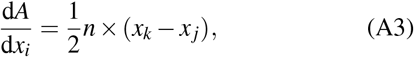

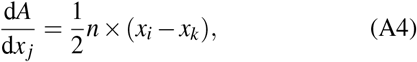

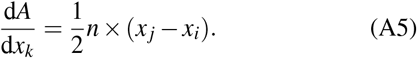

The gradient of the volume *V* associated to an oriented triangle (*i, j, k*) ∈ *F* (computed as the volume of the tetrahedron (*x*_*i*_, *x* _*j*_, *x*_*k*_, 𝒪), 𝒪 being the origin, is given by

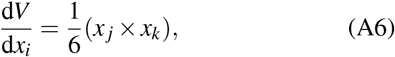

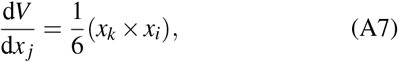

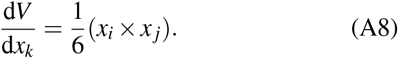

And the gradient of the dihedral angle *ϕ*_*i j*_ between the normals of the two oriented triangles (*i, j, k*) ∈ *F* and (*i, j, l*) ∈ *F*, with common edge (*i, j*) and normals *n*_1_ and *n*_2_, is given by

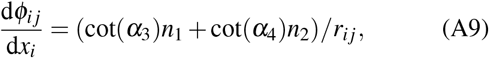

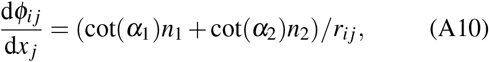

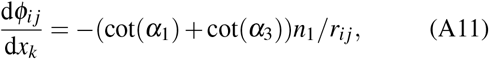

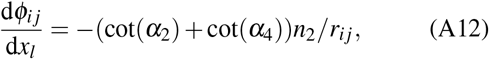

whereby the *α*_*i*_ are the sector angles defined by

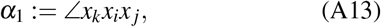

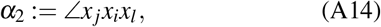

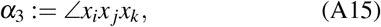

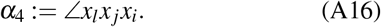

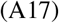

## Notes

https://github.com/bio-phys/trimem

### Competing Interest Statement

The authors have declared no competing interest.

