## Supplemental Figures for "TriMem: A Parallelized Hybrid Monte Carlo Software for Efficient Simulations of Lipid Membranes"

---

This document contains

- Supplementary Figures

---

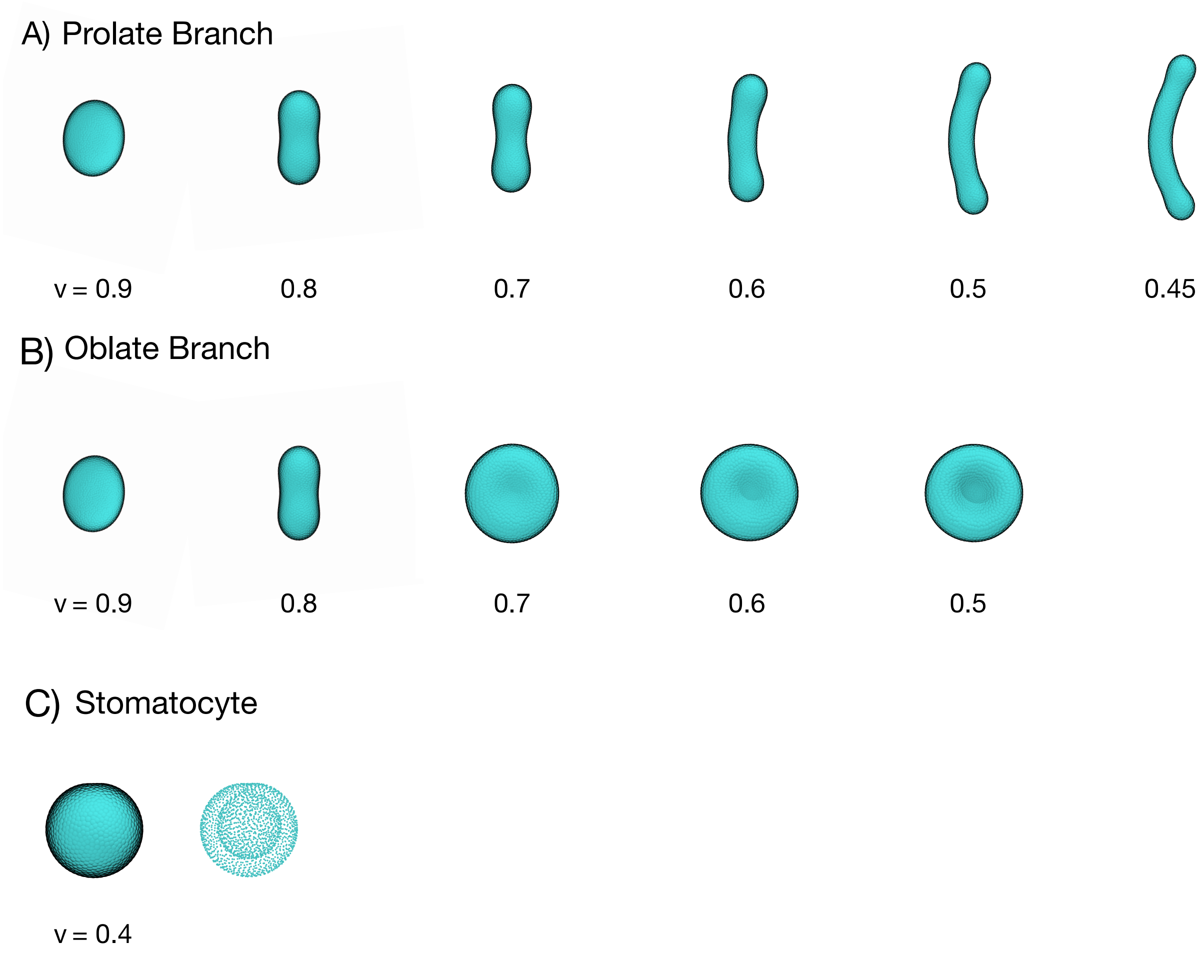


Figure S1: Minimized shapes with $N_V = 1962$ for the phase diagram of varying reduced volume $v = \{0.9, 0.8, 0.7, 0.6, 0.5, 0.45\}$, corresponding to Fig 6. (A) Shapes resulting from simulations initiated on the prolate branch. (B) Shapes resulting from simulations initiated from spheres/prolates. Note that for reduced volumes $v \geq 0.8$, oblate shapes spontaneously transitioned to prolate shapes. (C) Shape of an exemplary stomatocyte at volume $v = 0.4$. For better visibility the same structure is shown with smaller spheres on the right to reveal the inner shape. We note that the neck connecting the two spheres is very narrow. Vertices are shown as large cyan spheres.

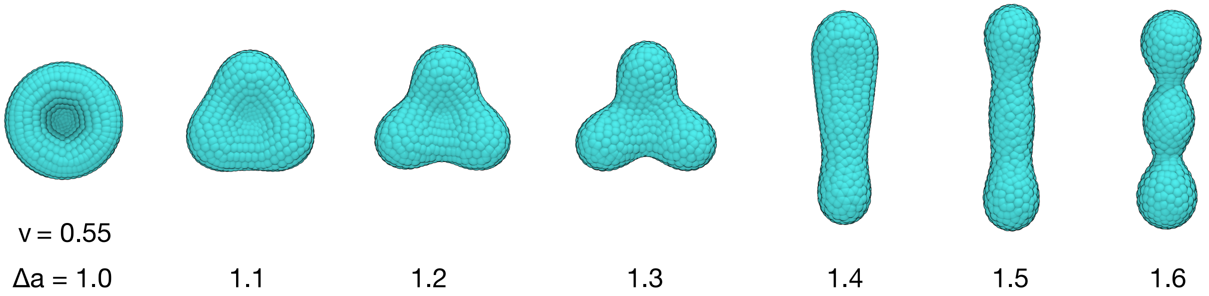


Figure S2: Minimized shapes with $N_V = 642$ for the phase diagram of varying area difference at reduced volume $v = 0.55$, varying area difference $\Delta A_0$ corresponding to Fig 7. Vertices are shown as large cyan spheres.
